# The aging rhythm: spatio-temporal dynamics of resting alpha oscillations in young and older brains

**DOI:** 10.64898/2026.07.30.741790

**Authors:** Andrea Alamia, Luca Tarasi, Pierre-Marie Matta, Jakob C. B. Schwenk, Vincenzo Romei

## Abstract

Aging is associated with substantial alterations in brain oscillatory activity, particularly within the alpha band (8–12 Hz). Yet, little is known about how aging affects the spatial propagation of alpha oscillations across cortical networks. In addition, although previous EEG studies have consistently reported age-related slowing of alpha peak frequency and changes in alpha power, the interpretation of these findings remains debated because oscillatory measures are influenced by age-related modifications in the aperiodic component of the power spectrum. Here, we investigated age-related changes in both the spectral and spatiotemporal properties of alpha activity using resting-state EEG data from a large cohort of younger (N = 326) and older adults (N = 108). To address the debate in the literature, analyses explicitly accounted for the aperiodic component of the EEG power spectrum. Consistent with previous literature, older adults exhibited a robust slowing of the individual alpha peak frequency, along with reductions in the aperiodic exponent and offset. Importantly, alpha-band power was also significantly reduced in older adults even after correcting for aperiodic activity, indicating that age-related alpha alterations cannot be fully explained by non-oscillatory spectral changes alone. Beyond conventional spectral measures, we characterized alpha-band traveling waves and identified age-related alterations in their propagation dynamics, particularly within frontal regions. Older adults showed enhanced medial-to-lateral and interhemispheric propagation patterns, suggesting reduced hemispheric segregation and increased bilateral coordination of rhythmic activity. These findings extend current models of cognitive aging by demonstrating that aging affects not only the spectral characteristics of alpha oscillations but also their large-scale spatiotemporal organization. Together, the results support the view that aging involves a functional reorganization of cortical communication dynamics, potentially reflecting compensatory mechanisms within distributed neural networks.

## Introduction

Brain oscillations support a wide range of cognitive functions by coordinating neuronal activity across distinct temporal and spatial scales (Ward, 2003; Buzsáki and Draguhn, 2004). Among these, alpha-band oscillations (8–12 Hz) are the most prominent rhythms observed in scalp electrophysiological (EEG) recordings in humans, and are widely distributed across cortical regions. Their functional significance has been extensively investigated across multiple cognitive domains (Palva and Palva, 2007; Klimesch, 2012), revealing their involvement in several complementary processes. Alpha oscillations have been linked to top-down mechanisms involved in inhibitory control, temporal coordination of cortical processing, and modulation of cortical excitability (Klimesch et al., 2007a; Jensen and Mazaheri, 2010; Sadaghiani and Kleinschmidt, 2016): changes in alpha power across occipital hemispheres reliably track shifts in visual attention, with increased power ipsilateral to the attended hemifield and decreased power contralaterally (Worden et al., 2000; Sauseng et al., 2005; Thut et al., 2006; Händel et al., 2011). Experimental evidence further shows that alpha oscillations can influence visual perception (Busch et al., 2009; Fakche et al., 2022; VanRullen and Macdonald, 2012; Romei and Tarasi, 2026): alpha activity in occipital and parietal regions has also been associated with perceptual processing and visual memory (Bonnefond and Jensen, 2012; Pang (庞兆阳) et al., 2020; Zeng et al., 2024; Wianda and Ross, 2019), revealing mechanisms that appear to reflect sensory processes distinct from alpha power changes linked to inhibitory control (VanRullen, 2016; Brüers and VanRullen, 2018; Keitel et al., 2019; Ruzzoli et al., 2019). Recent research on neural oscillations has increasingly focused on their spatial dynamics, particularly their propagation across cortical regions as traveling waves (TWs) (Muller et al., 2018). In scalp electrophysiological (EEG) recordings, TW propagation has been linked to several cognitive processes, including visual perception (Sato et al., 2012; Lozano-Soldevilla and VanRullen, 2019; Pang (庞兆阳) et al., 2020; Alamia et al., 2023; Tarasi et al., 2025a), working memory (Mohan et al., 2024; Zeng et al., 2024), and motor excitability (Zich et al., 2023; Haigh et al., 2025).

A growing body of research has demonstrated that aging is associated with substantial changes in brain oscillatory activity across multiple frequency bands (Dushanova and Christov, 2014; Ishii et al., 2017). Among these, alpha-band oscillations have received particular attention, as they undergo marked age-related alterations in both spectral power and peak frequency. Specifically, older adults typically exhibit a slowing of the individual alpha peak frequency (Klimesch, 1997; Chiang et al., 2011; Grandy et al., 2013; Stacey et al., 2021; Park et al., 2024), together with changes in alpha power distribution (Babiloni et al., 2006; Gómez et al., 2013; Ishii et al., 2017; Tröndle et al., 2023), reflecting modifications in large-scale neural dynamics and cortical processing efficiency. These alterations have been linked to age-related differences in sensory processing, attention, and memory performance, suggesting that alpha activity constitutes a sensitive marker of cognitive aging. However, the interpretation of age-related changes in alpha power remains debated, due to methodological confounds, such as variability in sample sizes and the influence of aperiodic neural activity on EEG power spectra. When accounting for this latter effect, some authors reported no differences in alpha power during aging (Scally et al., 2018; Merkin et al., 2023), suggesting that reported age-related reductions in alpha power may partly reflect changes in aperiodic activity rather than true oscillatory differences. In addition, while previous studies have extensively characterized the temporal and spectral properties of alpha oscillations in aging, considerably less attention has been devoted to their spatial dynamics. In particular, little is known about how aging affects the propagation of alpha-band oscillatory activity across cortical regions as traveling waves. Given the growing evidence linking traveling waves to perceptual and cognitive processes (Zhang et al., 2018; Lozano-Soldevilla and VanRullen, 2019; Alamia and VanRullen, 2023; Mohan et al., 2024), understanding whether and how their spatiotemporal organization changes with age may provide important insights into the neural mechanisms underlying cognitive aging.

Here, we addressed these limitations by analyzing a large resting-state EEG dataset (N=108 for the older group and N=326 for the younger group) while explicitly accounting for the aperiodic component of the power spectrum. This approach allowed us to reassess age-related changes in alpha oscillations with greater specificity. Consistent with previous findings, we observed a robust decrease in individual alpha peak frequency with aging. In contrast, although some recent studies reported no significant age-related differences in alpha power after separating periodic and aperiodic activity, we still identified reduced alpha power in older adults, suggesting that changes in aperiodic neural dynamics alone cannot fully explain age-related alterations in oscillatory activity. Importantly, beyond conventional spectral measures, our study provides a novel characterization of age-related changes in alpha-band traveling waves. We show that aging is associated with specific alterations in the spatial propagation of alpha activity, particularly within frontal regions. These findings extend current models of cognitive aging by highlighting that aging affects not only the spectral properties of alpha oscillations, but also their large-scale spatiotemporal organization across cortical networks.

## Material and Methods

### Participants

Participants were pooled from multiple datasets with distinct experimental designs (Ursino et al., 2022; Tarasi et al., 2025b, 2025c; Trajkovic et al., 2025a; Tarasi et al., 2026b), sharing the same EEG setup. In each dataset, individuals completed one to two minutes of resting-state recording under both eyes-closed and eyes-open conditions prior to task performance. The present study focuses exclusively on these resting-state segments. The final sample comprised four groups: 326 younger participants in the eyes-closed condition (mean age = 23.17 ± 3.11 years, range = 18–43; 203 women) and 235 younger participants in the eyes-open condition (mean age = 22.97 ± 2.81 years, range = 18–35; 150 women), as well as 108 older participants in the eyes-closed condition (mean age = 64.16 ± 7.56 years, range = 55–88; 59 women) and 63 older participants in the eyes-open condition (mean age = 66.50 ± 8.41 years, range = 57–87; 37 women). Chi-square tests confirmed no significant difference in gender distribution between younger and older participants in either the eyes-closed condition (chi(1) = 1.98, p = .159) or the eyes-open condition (chi(1) = 0.55, p = .457), nor between conditions within the younger (chi(1) = 0.14, p = .706) or older group (chi(1) = 0.27, p = .602).

### EEG preprocessing

64-electrode setup was mounted according to the International 10–10 system. EEG was measured with respect to FCz electrodes, with all electrode impedances maintained below 10 kΩ. EEG signals were acquired at a rate of 1000 Hz and processed offline using custom MATLAB scripts (Version R2022b) in combination with the EEGLAB toolbox (Delorme & Makeig, 2004). The EEG recording was filtered offline in the 0.5- to 70-Hz band and a notch filter (50Hz) was applied. Visual inspection was performed to identify noisy channels which were subsequently spherically interpolated, and recording segments corrupted by artifacts were eliminated. The recording was then referenced to the average of all electrodes. Independent Component Analysis (ICA) was applied to identify and remove artifacts that were clearly distinguishable from brain-origin EEG signals.

### Oscillatory power and FOOOF analysis

Power spectral density (PSD) estimates were computed using the *ft_freqanalysis* function from the FieldTrip toolbox. Continuous EEG data were downsampled from 1000Hz to 100Hz and transformed into the frequency domain using a Fourier-based approach (i.e., multitaper with the Hanning method). To separate periodic (oscillatory) and aperiodic (1/f-like) components of the spectra, the FOOOF (Fitting Oscillations and One-Over-F) algorithm was applied to the resulting PSDs (Donoghue et al., 2020). The FOOOF model parameterizes the aperiodic component as a broadband offset and slope, which are estimated for each electrode before removal from the spectra to isolate oscillatory activity. Removal was performed by computing the quotient between the aperiodic component and the original spectra. Model fitting was performed using standard FOOOF settings. The resulting aperiodic-adjusted spectra were used for all electrodes to estimate spectral power in the alpha band, defined as 8-12 Hz. In addition, we compared spectral power between groups within the individual alpha frequency (IAF) ± 2 Hz, to account for differences in the alpha range between groups. The alpha peak was identified within the range 5-15Hz and discarded if smaller than 7 Hz or larger than 13 Hz. Overall, one participant (older adult, eyes-open condition) was excluded because no identifiable peak could be detected in the alpha range over occipital electrodes (all three posterior regions, as defined below). Group differences were first assessed across all electrodes and subsequently within six predefined regions of interest (ROIs), following a previous approach (Merkin et al., 2023). The ROIs comprised anterior left (AF3, F3, F5, F7, FC3, FC5, FT7, C3, C5, T7), anterior midline (Fz, FC1, FC2, F1, F2, C1, C2), anterior right (AF4, F4, F6, F8, FC4, FC6, FT8, C4, C6, T8), posterior left (TP7, CP3, CP5, P3, P5, P7, PO3, PO7, O1), posterior midline (Cz, CPz, CP1, CP2, Pz, P1, P2, POz, Oz), and posterior right (TP8, CP4, CP6, P4, P6, P8, PO4, PO8, O2).

### Traveling waves analysis

Traveling waves were quantified using two complementary approaches: a spectral method based on a two-dimensional Fast Fourier Transform (2D-FFT) (Alamia and VanRullen, 2019; Wei et al., 2024; Zeng et al., 2024) and a spatial phase-gradient method based on linear plane fitting. Both methods were applied to characterize the direction and strength of wave propagation across the scalp (Zhang et al., 2018; Das et al., 2022; Schwenk and Alamia, 2025).

#### 2D-FFT analysis

For the 2D-FFT analysis, 1-s segments were extracted from midline electrodes (Oz, POz, Pz, CPz, Cz, FCz, and Fz) using a sliding window with 500-ms overlap. Signals were arranged into two-dimensional matrices with electrodes along the spatial axis and time along the temporal axis. A 2D-FFT was applied to each matrix to obtain the joint spatial–temporal frequency spectrum. In the resulting spectra, the upper and lower quadrants correspond to forward (FW) and backward (BW) propagating waves as a function of temporal frequency. Wave power was quantified by averaging spectral power within the alpha band (8–12 Hz). Values were expressed in decibels as the logarithmic ratio between these measures and the average temporal power obtained by applying a one-dimensional FFT (1D-FFT) independently to each electrode. Because the 2D-FFT captures both spatial propagation and the temporal spectral structure of individual signals, a surrogate baseline was computed to isolate the spatial propagation component. Specifically, electrode order was randomly shuffled before constructing the spatiotemporal matrices, thereby disrupting spatial phase relationships while preserving each signal’s temporal properties. The full analysis was repeated on the shuffled data. Traveling wave power was then quantified by comparing ordered and shuffled spectra using a logarithmic ratio across frequencies between 2 and 45 Hz. Following previous studies (Alamia and VanRullen, 2019; Pang (庞兆阳) et al., 2020), the maximum values in the 2D-FFT spectra for the real data (FW and BW) were divided by the corresponding values obtained from shuffled data (FWss and BWss), yielding wave power expressed in decibels (dB).

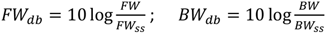

#### Phase fitting analysis

Traveling waves were additionally characterized using a spatial phase-gradient approach based on linear plane fitting of instantaneous phases. Continuous phase time series were computed for electrodes within a predefined region of interest (ROI) comprising frontal and parieto-occipital electrodes (occipital electrodes: O1, Oz, O2, PO7, PO3, POz, PO4, PO8, P7, P5, P3, P1, Pz, P2, P4, P6, P8; frontal electrodes: Fp1, Fp2, AF7, AF5, AF3, AFz, AF2, AF4, AF8, F7, F5, F3, F1, Fz, F2, F4, F6, F8). Analyses were performed in the alpha band (8–12 Hz). Signals were band-pass filtered using a finite impulse response (FIR) filter, and instantaneous phases were extracted via the Hilbert transform. Phases were referenced to the mean phase within the ROI. Electrode positions were projected onto a two-dimensional plane using the standard 10–20 layout implemented in the FieldTrip toolbox (Oostenveld et al., 2011). For each time point, spatial phase gradients were estimated by fitting a linear plane to the phase distribution across electrodes:

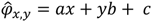

where *x* and *y* denote electrode coordinates, *a* and *b* represent spatial phase gradients along the two axes, and *c* corresponds to a constant phase offset. The propagation direction of the wave was defined by the orientation of the phase gradient:

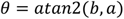

For each time point, plane fits were evaluated across possible propagation directions, and the optimal solution was selected by maximizing the vector length of residuals in circular phase space. The goodness of fit was quantified by the proportion of variance explained (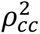), computed from the circular correlation between predicted and observed phases. A surrogate null distribution of 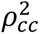 values was generated using a bootstrap procedure in which electrode positions were randomly permuted 10 times, yielding a subject-specific distribution. Plane fits obtained from the real data were retained only if their 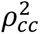 values exceeded the 99^th^ percentile of the corresponding null distribution. For each condition and time point, the proportion of trials exhibiting forward (FW) and backward (BW) propagating waves was then computed. Propagation direction was defined based on the estimated gradient angle α, with FW and BW waves corresponding to α=π and α=0 radians, respectively, using a tolerance of ±0.5 radians. The retained plane fits also yielded estimates of the spatial frequency. Combined with the temporal frequency, these estimates were used to calculate traveling-wave propagation velocity as the ratio of temporal to spatial frequency. Propagation velocity was estimated for alpha-band traveling waves in younger and older adults during both eyes-open and eyes-closed conditions.

### Statistical analysis

Since we have two groups with an unequal number of participants and conditions, we performed all statistical analyses using Linear Mixed Models. Degrees of freedom were estimated using the Satterthwaite approximation. In addition, Bayesian ANOVA was performed to obtain Bayes factors (BFs) for each effect of interest, specifically to estimate effect sizes and corroborate the absence of an effect. Bayes factors quantify the relative evidence for the alternative hypothesis compared with the null hypothesis. Throughout the manuscript, we report both p-values from the linear Mixed models and the Bayes factors in favor of the alternative hypothesis (BF₁₀). All models included the CONDITION (closed or open eyes) as a within-subject factor, the GROUP (older or younger) as a between-subject factor, and their interaction. PARTICIPANT was considered a random term. All statistical analyses were conducted in JASP (Love et al., 2019) using the default prior specifications. The association between traveling-wave speed and individualized alpha frequency was assessed using Spearman’s rank correlation coefficient, which does not assume linearity or normality.

## Results

### Aperiodic and periodic spectral components

We first assessed differences in the aperiodic component between the two age groups by comparing the slope and intercept estimated by the FOOOF method (see methods for details). As shown in Figure 1, we observed a difference between the two groups both in the intercept (F(1,718.39)=136.5, p<0.001, BF_10_>10^14^) and in the slope (F(1,713.06)=74.73, p<0.001, BF_10_>10^14^), as well as a difference between the two conditions ‘closed eyes’ and ‘open eyes’ (effect of condition in the intercept: F(1,560.8)=46 p<0.001, BF_10_>10^13^ and in the slope: F(1,555.7)=20.6, p<0.001, BF_10_>10^9^).In addition, we also found a significant interaction between GROUP and CONDITION in the slope (F(1,555.7)=5.07, p=0.025, BF_10_=8.49) but not in the intercept (F(1,560.8)=0.153, p=0.695, BF_10_=0.58). Figure 1 illustrates the spatial distribution of both parameters across conditions for the two groups. Following previous work (Merkin et al., 2023), we assessed regional differences by grouping electrodes into six regions: anterior-left, anterior-midline, anterior-right, posterior-left, posterior-midline, and posterior-right. We then repeated the analyses described above, including **REGION** as an additional factor. As shown in Supplementary Figure S1, we observed effects similar to those reported above for both the intercept and slope. Significant main effects were found for GROUP (intercept: F(1,126.37)=152.6, p<0.001, BF_10_>10^12^, slope F(1,111.65)=85.68, p<0.001, BF_10_>10^9^) and CONDITION (intercept: F(1,123.58)=38.68, p<0.001, BF_10_>10^11^, slope F(1,128.47)=10.57, p<0.001, BF_10_>10^7^). The **GROUP × CONDITION** interaction was significant for the slope (F(1,102.21)=4.62, p=0.034, BF_10_>10^6^) but not for the intercept (F(1,103.65)=0.031, p=0.858, BF_10_>10^6^). A significant main effect of **REGION** was observed for both the intercept (F(5,657.88)=265.7, p<0.001, BF_10_>10^17)^, and slope (F(5,994.57)=114.7, p<0.001, BF_10_>10^15^In addition, all two-way and the three-way interactions involving **REGION** were significant (all p<0.001, and BF_10_>10^7^). As shown in figure 1, the younger group exhibited higher intercept values in central regions and along the midline, along with steeper slopes in frontal and occipital areas. In contrast, the older group exhibits lower overall values and a topography that is less focal and more diffusely distributed.

**Figure 1:**
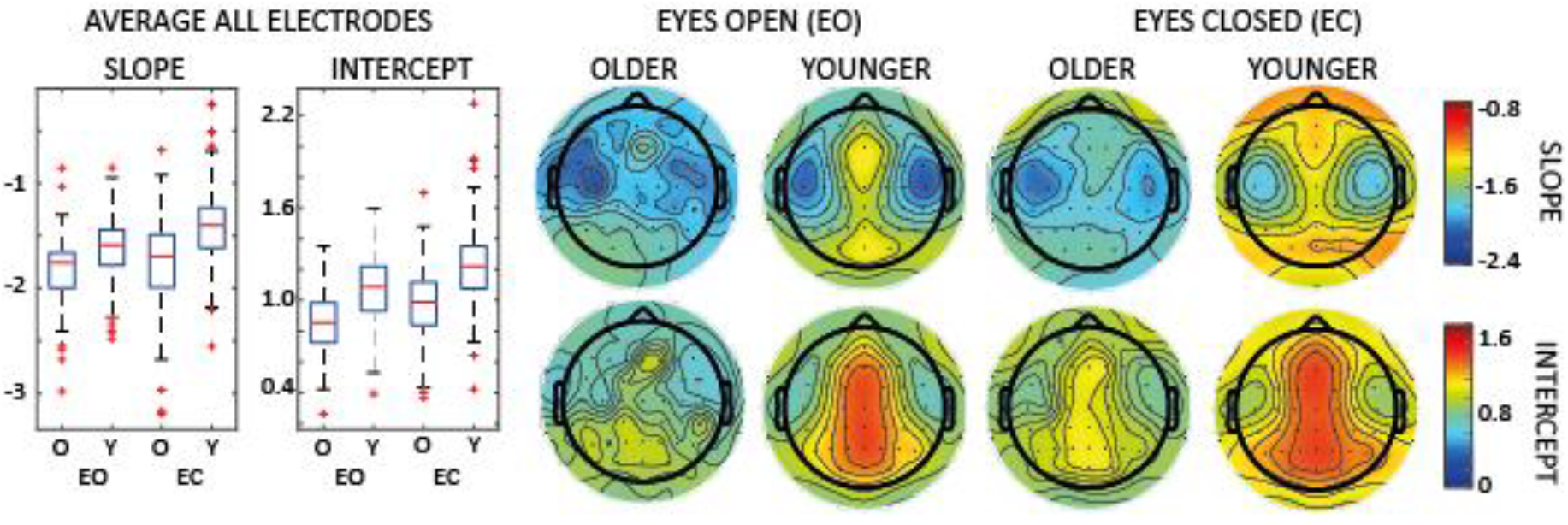
Group differences in aperiodic spectral parameters during resting-state EEG. Boxplots show the average values across all electrodes for the aperiodic **slope** (left) and **intercept** (middle) in older (O) and younger (Y) adults during eyes-open (EO) and eyes-closed (EC) resting-state recordings. Boxes indicate the interquartile range (IQR), center lines denote the median, whiskers extend to 1.5 × IQR, and red dots represent individual participants. Scalp topographies show the spatial distribution of the aperiodic slope (top row) and intercept (bottom row) for younger and older adults during EO and EC. Warmer colors indicate higher parameter values and cooler colors indicate lower parameter values. Contour lines represent interpolated isovalues across the scalp. Topographic maps are displayed with the nose at the top and the left hemisphere on the left.

### Alpha-band topography, power, and frequency in the younger and elderly populations

After quantifying the difference between groups in the aperiodic component, we investigated differences in the periodic component, specifically in the alpha band. In particular, we quantified the oscillatory power and peak frequency after removing the aperiodic component. Statistical analysis revealed a large difference between groups in both frequency (F(1,726)=34.1, p<0.001, BF_10_>10^7^) and power (F(1,687.7)=17.7, p<0.001, BF_10_>10^3^). As illustrated in Figure 2, we observed stronger alpha power in the younger group, specifically in occipital regions, and, as expected, greater power in both groups during the eyes-closed condition (effect of condition: F(1,512.9)=87, p<0.001, BF_10_>10^14^). We also observed a higher alpha-band frequency in the younger group, but no differences between eyes-open and eyes-closed conditions (F(1,726)=0.765, p=0.382, BF_10_=0.084). Lastly, we did not find any significant interaction between GROUP and CONDITION (power: F(1,512.9)=2.7, p=0.101, BF_10_=2.358; frequency: F(1,726)=0.30, p=0.58, BF_10_=0.072). As for the aperiodic analysis, we included the factor REGION to assess differences in different areas (Merkin et al., 2023), obtaining consistent results. As shown in Supplementary Figure S2, we found significant main effects for GROUP (power: F(1,222.05)=55.37, p<0.001, BF_10_>10^13^, α-frequency F(1,186.57)=12.03, p<0.001, BF_10_>10^12^) and CONDITION (power: F(1,243.29)=164.3, p<0.001, BF_10_>10^13^, α-frequency F(1,366.82)=84.68, p<0.001, BF_10_>10^12^). The **GROUP × CONDITION** interaction was significant for the α-frequency (F(1,2973,30)=23.35, p=0.034, BF_10_>10^5^) and for the power (F(1,203.86)=0.031, p=0.039, BF_10_=1315). A significant main effect of **REGION** was observed for both the power (F(5,720.70)=74.11, p<0.001, BF_10_>10^13)^, and α-frequency (F(5,693.83)=38.49, p<0.001, BF_10_>10^15^). In addition, all two-way and three-way interactions involving **REGION** were significant (all p<0.001, and BF_10_>10^7^). Figure 2 shows that the younger group exhibited higher alpha-band power in occipital and frontal regions, together with higher peak alpha frequencies in parietal and occipital areas. By contrast, the older group showed lower alpha-band values overall and a less focal, more spatially diffuse topographical distribution. As a robustness analysis, we quantified alpha-band power within an individualized frequency window centered on each participant’s individualized alpha frequency (IAF ± 2 Hz) to account for the age-related slowing of alpha oscillations. The results were consistent with those obtained using the fixed alpha band, showing significant main effects of group (F(1,474.21)=31.23, p<0.001, BF_10_=6591) and condition (F(1,402.24)=84.42, p<0.001, BF_10_>10^14^), together with a significant group × condition interaction (F(1,476.22)=3.852, p=0.050, BF_10_=2.251; Supplementary Fig. S3).

**Figure 2:**
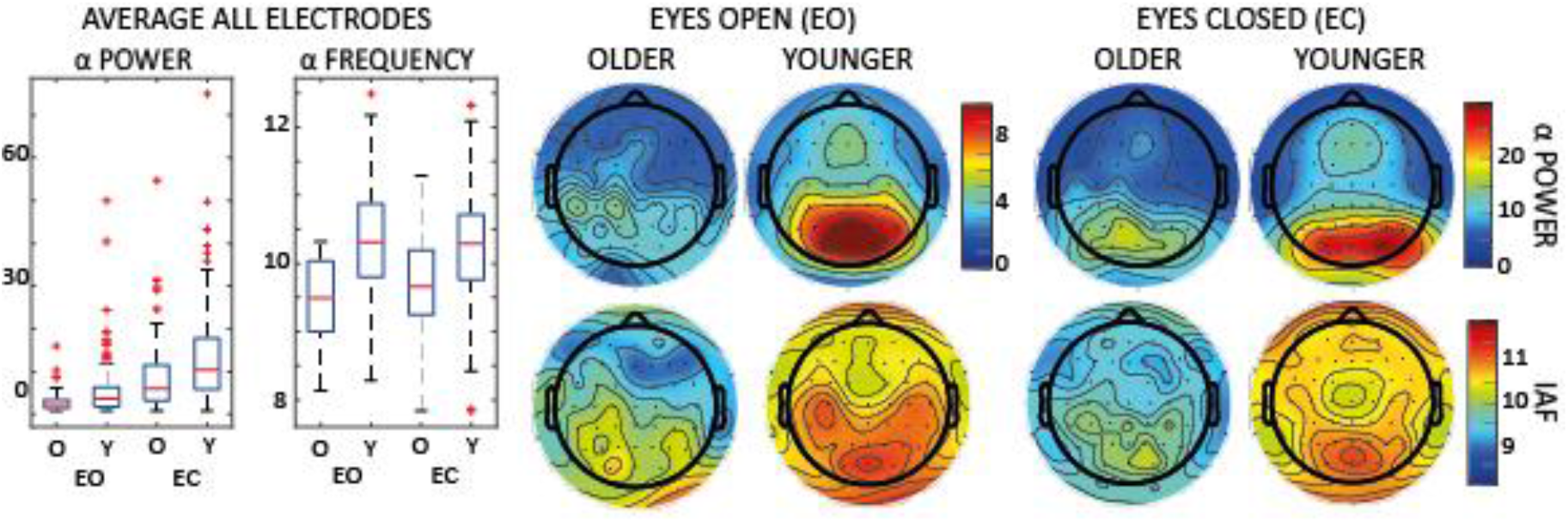
Group differences in alpha oscillatory activity during resting-state EEG. Boxplots show the average values across all electrodes for **alpha power** (left) and **individual alpha frequency (IAF)** (middle) in older (O) and younger (Y) adults during eyes-open (EO) and eyes-closed (EC) resting-state recordings. Boxes indicate the interquartile range (IQR), center lines denote the median, whiskers extend to 1.5 × IQR, and red dots represent individual participants. Scalp topographies show the spatial distribution of alpha power (top row) and PAF (bottom row) for younger and older adults during EO and EC. Warmer colors indicate higher values and cooler colors indicate lower values. Contour lines represent interpolated isovalues across the scalp. Topographic maps are displayed with the nose at the top and the left hemisphere on the left.

### Spectra of forward and backward traveling waves

We then characterize the amount of oscillations propagating as traveling waves in the forward (FW) or backward (BW) direction along the midline of electrodes (i.e., the Oz–Fz axis) and as a function of their temporal frequency, using a method based on the 2D Fast Fourier Transformation (Pang (庞兆阳) et al., 2020; Zeng et al., 2024; Luo and Ester, 2025). Figure 3 illustrates the spectral profiles of FW and BW waves for both groups: interestingly, we found that alpha- and low-beta-band oscillatory waves propagate mostly in the top-down direction as BW waves, whereas high-beta activity (above 25Hz) propagates in the forward direction. These results align with previous studies linking higher-frequency bands (i.e., high-beta/gamma) to forward processing and lower frequencies to top-down processing (Bauer et al., 2014; Bastos et al., 2015; Bressler and Richter, 2015; Michalareas et al., 2016; Trajkovic et al., 2025b). Regarding alpha-band oscillations, we observed a remarkable difference between the two groups in the forward waves (F(1,728)=30.68, p<0.001, BF_10_>10^6^), but not in the backward waves (F(1,728)=0.64, p=0.424, BF_10_=0.077), and we did not report any difference between eyes open and closed conditions (forward: F(1,728)=0.11, p=0.741, BF_10_=0.08; backward: F(1,728)=0.005, p=0.943, BF_10_=0.09) or the interaction (forward: F(1,728)=0.22, p=0.635, BF_10_=0.06; backward: F(1,728)=1.911, p=0.167, BF_10_=0.18).

**Figure 3:**
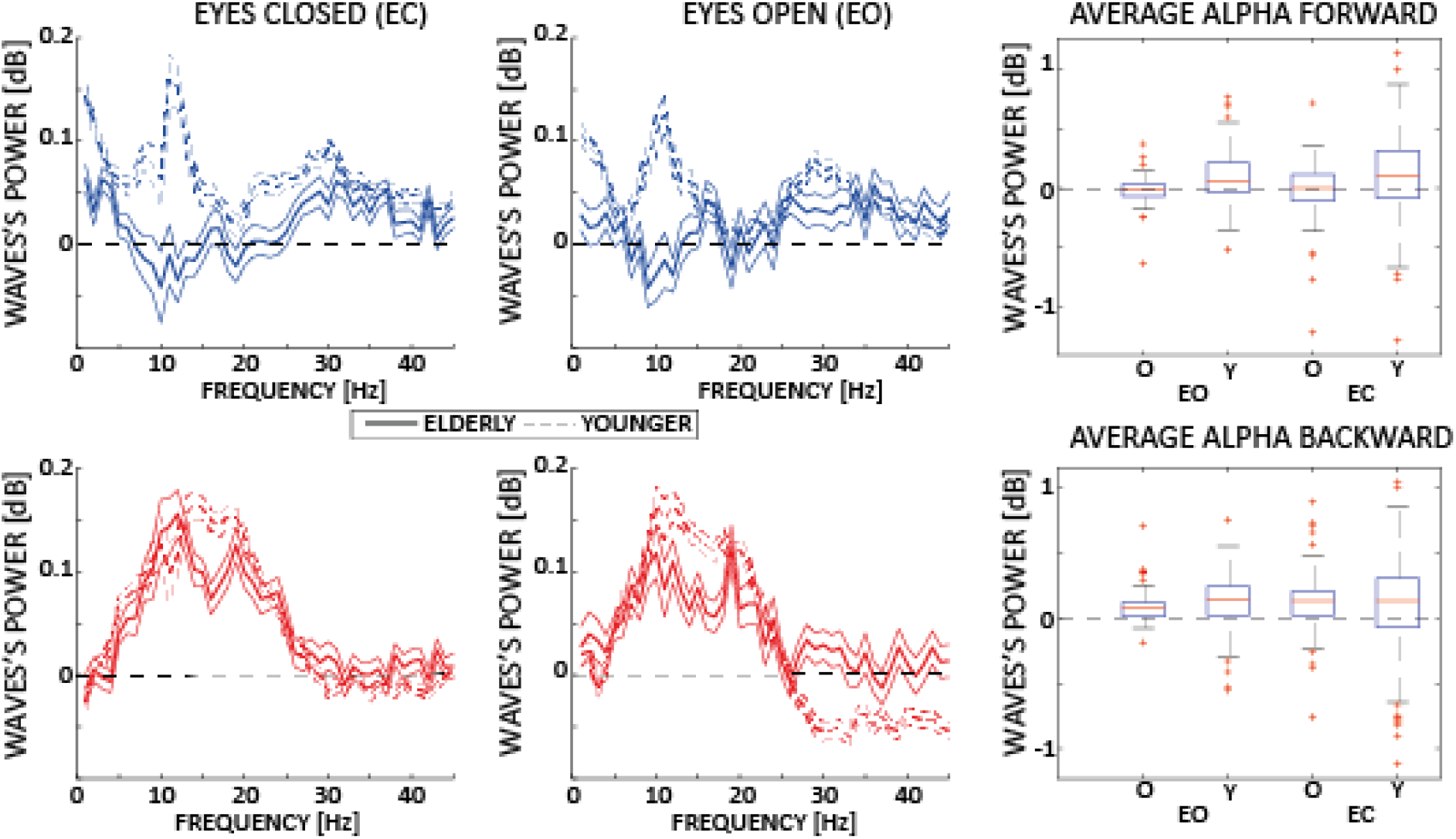
Spectra of traveling waves during eye closed and open for younger and older adult groups. Average and standard error of the mean of the amount of waves in dB computed for the younger (dashed) and older (solid lines) populations for both forward (in blue) and backward (in red) waves along the midline of electrodes. The boxplots represent the amount of alpha-band waves under both resting-state conditions. Boxes indicate the interquartile range (IQR), center lines denote the median, whiskers extend to 1.5 × IQR, and red dots represent individual participants. Overall, younger populations have stronger forward waves than older populations.

### Disentangling forward and backward traveling waves in occipital and frontal regions

We further investigated differences in the spatiotemporal organization of traveling waves between the two groups by complementing the 2DFFT approach with a phase-gradient–based method (Zhang et al., 2018; Schwenk and Alamia, 2025). This approach offers the advantage of quantifying wave propagation within specific regions of interest while being less influenced by oscillatory power. Specifically, it estimates propagation direction by fitting a spatial phase gradient on a spherical model within a defined frequency band (i.e., alpha band) and across two regions of interest: frontal and occipital electrodes (Figure 4). As illustrated in Figure 4, linear mixed-effects modeling revealed a comparable pattern of traveling waves between groups in occipital regions. There was no significant effect of GROUP (F(1,7701)=0.776, p=0.379), CONDITION (F(1,6048)=0.168, p=0.681), or their interaction (F(1, 6048)=1.199, p=0.274), but a significant main effect of ANGLE (F(1, 3992)=18.318, p<0.001). In contrast, a different pattern emerged in frontal electrodes. Here, we observed significant effects of GROUP (F(1, 7855)=57.583, p<0.001), ANGLE (F(1, 3696)=50.533, p<0.001), and their interaction (F(1, 7855)=102.539, p<0.001), whereas CONDITION was not significant (F(1, 6126)=0.878, p=0.349). These findings indicate a region-specific alteration in wave propagation: in younger participants, alpha waves predominantly propagate along the fronto-occipital axis, whereas in older participants they preferentially travel along the orthogonal medio-lateral axis. Finally, we observed significant group differences in the speed of traveling waves in both occipital and frontal regions, having faster traveling waves in the younger group than in the elder one in both conditions (Figure 4). This was supported by main effects of GROUP (occipital: F(1, 409.878)=58.409, p<0.001, BF10>102; frontal: F(1, 4486.414)=83.869, p<0.001, BF10>1014) and CONDITION (occipital: F(1, 25.348)=3.612, p=0.05, BF10>1014; frontal: F(1, 5279)=98.692, p<0.001, BF10>1014), with no significant GROUP × CONDITION interaction (occipital: F(1, 2.089)=0.298, p=0.586, BF10=0.710; frontal: F(1, 82.827)=1.548, p=0.214, BF10=1.041). Interestingly, traveling-wave speed was negatively correlated with individualized alpha frequency in the younger group, but not in the older group. In younger adults, significant associations were observed in posterior regions during the eyes-closed condition (left: ρ=-0.2066, p=0.0002; middle: ρ=-0.1475, p=0.0078; right: ρ=-0.2115, p=0.0001), and in frontal regions during the eyes-open condition (left: ρ=-0.1982, p=0.0023; middle: ρ=-0.1413, p=0.0319; right: ρ=-0.1485, p=0.0237). No significant correlations were observed in the older group (all P>0.1).

**Figure 4:**
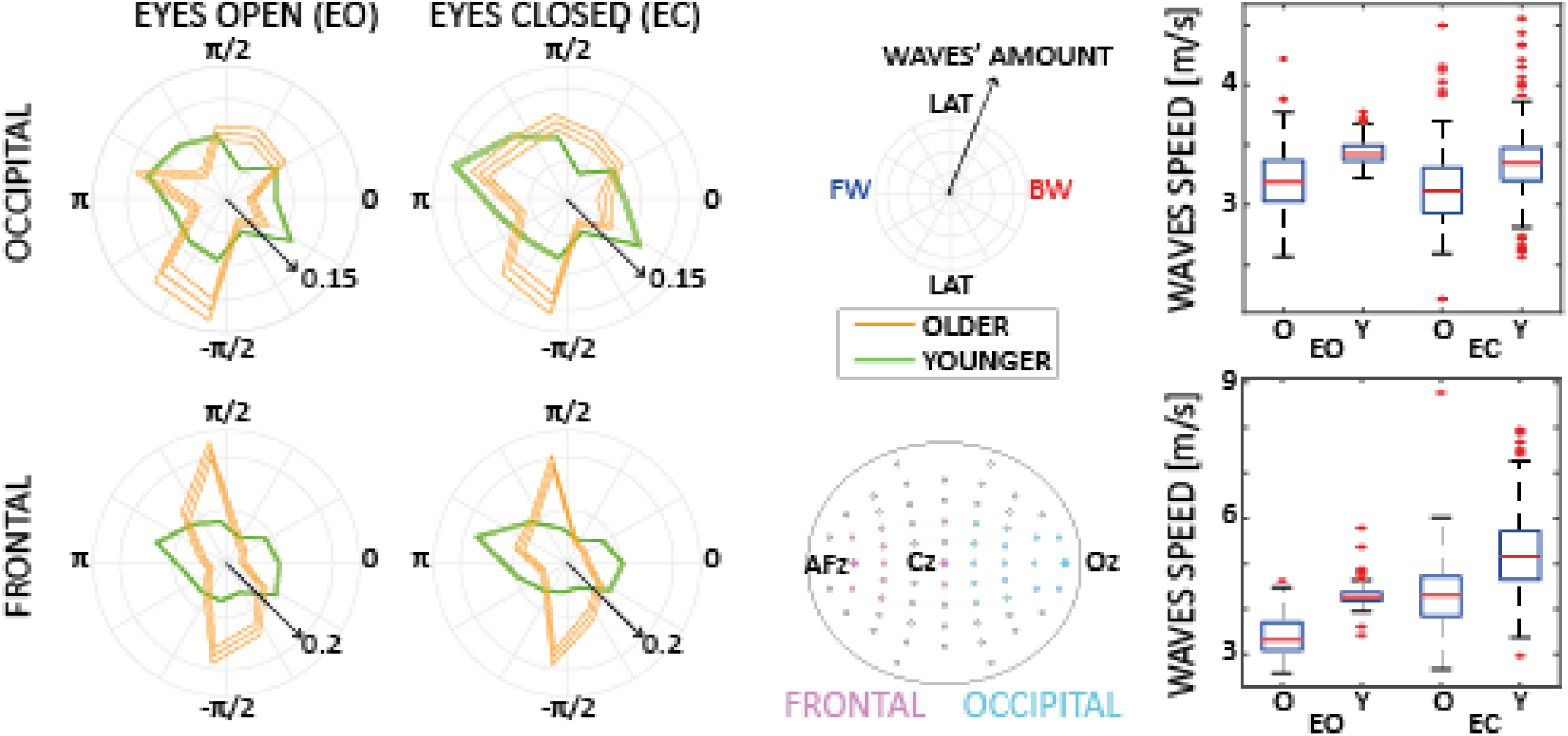
Traveling waves analysis in frontal and occipital regions. The polar plots show the average direction in frontal and occipital regions (electrodes showed in purple and cyan, respectively). Direction at 0 radians show backward waves (from frontal to occipital regions), whereas *π* radians reveal forward waves (from occipital to frontal). ±*π*/2 correspond to medial-to-lateral waves in the two hemispheres. The boxplots to the right show the average speed of the alpha-band waves in each region for the two populations in the two conditions. Interestingly, we observed a significantly different pattern in frontal regions between the two groups, with a significantly larger medio-lateral propagation in the elderly group, compared to a more posterior-anterior orientation in the younger group.

## Discussion

In this study, we characterize how aging impacts both the spectral and spatiotemporal properties of alpha-band activity in humans. Using a large EEG dataset and explicitly accounting for the aperiodic component of the power spectrum, we replicated the well-established age-related slowing of the individual alpha peak frequency, changes in the aperiodic component, and a reduction of alpha power in older adults. These results suggest that alterations in alpha oscillations with aging are unlikely to be fully attributed to changes in aperiodic neural activity alone. Beyond conventional spectral measures, we further demonstrate that aging is associated with specific changes in the propagation of alpha-band traveling waves, particularly across frontal regions, pointing to an age-related reorganization of large-scale cortical dynamics.

Our findings of reduced alpha-band power in older adults during eyes-closed resting state are broadly consistent with a substantial body of previous EEG aging literature. Earlier studies reported robust age-related reductions in alpha power during resting-state conditions (Polich, 1997; Babiloni et al., 2006), supporting the view that aging is associated with diminished oscillatory synchronization within canonical alpha networks. More recently, however, this interpretation has been refined by studies emphasizing the contribution of the aperiodic component of the EEG spectrum. For example, some authors argued that apparent reductions in alpha power with age may largely reflect changes in the underlying aperiodic background activity rather than true oscillatory alterations per se (Merkin et al., 2023). Nevertheless, our results aligned with findings suggesting that both alpha power and aperiodic parameters (offset and exponent) decrease with age, and importantly, that alpha reductions remain significant even after correcting for aperiodic activity (Tröndle et al., 2023). Together, these findings suggest that previously reported age-related changes in alpha activity likely reflect contributions from both periodic alpha oscillations and the aperiodic spectral background, which can be dissociated using spectral parameterization. The present findings also fit within the broader literature documenting the slowing of alpha rhythms with aging (Klimesch, 1999; Knyazeva et al., 2018). Several studies have consistently reported reductions in individual or peak alpha frequency in older adults. Previous work observed age-related slowing of individual alpha frequency (IAF), although without reductions in alpha power (Stacey et al., 2021). In contrast, a recent meta-analysis demonstrated converging evidence for both slower peak alpha frequency and reduced alpha power in elderly populations, with the strongest effects localized to occipital and sensorimotor/parietal regions (Park et al., 2024). All in all, our findings are more consistent with this latter account, supporting the notion that aging is accompanied by both spectral slowing and diminished alpha-band power. The alpha slowdown may also help explain age-related declines in visual performance and temporal sensitivity, as faster alpha rhythms have been associated with better visual sensitivity and temporal discrimination (Samaha and Romei, 2024; Tarasi and Romei, 2024; Frisoni et al., 2025; Romei and Tarasi, 2026). Accordingly, alpha slowing may contribute to less efficient visual and temporal processing in aging (He et al., 2020, 2025).

Age-related increases in interhemispheric connectivity in frontal regions observed in the present study are broadly consistent with previous models of functional reorganization in aging, suggesting a more integrated and less efficient architecture (Tomasi and Volkow, 2012; Reuter-Lorenz and Park, 2014; Spreng and Turner, 2019; Deery et al., 2023). Particularly, the HAROLD (Hemispheric Asymmetry Reduction in Older Adults) framework (Cabeza, 2002; Berlingeri et al., 2013) proposes that older adults exhibit reduced hemispheric lateralization during cognitive processing, often interpreted as reflecting either compensatory recruitment or neural dedifferentiation. However, HAROLD was originally derived from task-related activation patterns, primarily in episodic and working memory paradigms, leaving open the question of whether its core principles extend to intrinsic neural dynamics at rest. In our data, older adults demonstrated greater medial-to-lateral connectivity in alpha-band traveling waves in frontal regions, indicative of stronger interhemispheric interactions than in younger participants. This pattern is compatible with the overall reduction of 1) long-range connections (Damoiseaux, 2017), 2) reduction of alpha-band coherence (Vysata et al., 2014), and 3) reduction of hemispheric specialization in aging (Dolcos et al., 2002). Moreover, this pattern extends the HAROLD framework from regional activation patterns to the dynamics of large-scale oscillatory propagation. The findings may therefore suggest that aging is associated not only with bilateral recruitment of cortical regions, but also with increased coordination between hemispheres at the level of rhythmic network communication. Future studies on broadband dynamics will test this hypothesis more directly. The present results may also relate, albeit more indirectly, to the Posterior–Anterior Shift in Aging (PASA) model (Davis et al., 2008). PASA describes a characteristic reduction in posterior activity accompanied by increased frontal recruitment in older adults, often interpreted as compensatory support for preserved cognitive performance. The observed increase in interhemispheric alpha-band connectivity in frontal areas could reflect a specific reorganization of functional communication networks that accompanies compensatory processing in aging (Reuter-Lorenz and Cappell, 2008). In this sense, enhanced medial-to-lateral propagation may represent a complementary mechanism to the frontal recruitment emphasized in PASA, whereby older adults rely on more distributed and less segregated network dynamics (Deery et al., 2023). Future work relating traveling-wave connectivity measures directly to behavioral performance may help strengthen this interpretation and assess its functional relevance.

This functional reorganization of alpha-band propagation may also align with recent evidence showing that aging shifts the site at which prior expectations are integrated in perceptual decision-making, with older adults relying more heavily on fronto-central, action-related rather than parieto-occipital, perception-related processing (Tarasi et al., 2026b). Considering alpha oscillations as the neural substrate for prior expectations (Rohenkohl and Nobre, 2011; van Diepen et al., 2015; Mayer et al., 2016), the enhanced fronto-lateral and interhemispheric alpha propagation observed here may provide a candidate oscillatory substrate for such a shift: a redistribution of large-scale rhythmic communication away from posterior sensory hubs toward more anterior, bilaterally coordinated networks may underpin the increased weight of action-bound priors in older adults. While speculative, this convergence between resting-state oscillatory reorganization and task-based perceptual-motor shifts suggests that altered alpha traveling-wave dynamics may constitute a system-level signature of the broader functional remapping that characterizes the aging brain (Sala-Llonch et al., 2015; Damoiseaux, 2017).

Our findings are partially consistent with prior studies reporting age-related reductions in functional coupling within the alpha band (Ishii et al., 2017; Lejko et al., 2020). In particular, previous resting-state EEG investigations have shown that older adults exhibit significantly weakened global connectivity measures such as PLI and WPLI in the upper alpha range (Kikuchi et al., 2000; Gaál et al., 2010; Vecchio et al., 2014; Scally et al., 2018), suggesting a decline in the efficiency or stability of long-range oscillatory communication with aging. At first glance, these findings may appear in contrast with our observation of increased medial-to-lateral and interhemispheric connectivity in alpha-band traveling waves. However, these results are not necessarily contradictory, as they may reflect different aspects of network organization. The overall reduction in global synchronization efficiency is not incompatible with a redistribution toward more diffuse or bilateral propagation dynamics. As discussed above, such a pattern would be compatible with theories of reduced functional specialization and increased compensatory recruitment in older adults. More broadly, our results support the notion that aging is accompanied by large-scale reorganization of resting-state brain dynamics (Mccarthy et al., 2014; Li et al., 2015). Previous studies demonstrated an age-related anterior shift in oscillatory activity during rest, extending the PASA-like posterior-to-anterior redistribution beyond task-related paradigms (Perinelli et al., 2022). Connectivity analyses further suggested that aging alters the spatial organization of resting-state networks rather than simply reducing overall activity (Balsters et al., 2013; Song et al., 2014). In parallel, studies examining signal complexity have reported that measures such as fractal dimension decline in older adults following increases during earlier adulthood, indicating reduced complexity and flexibility of neuronal dynamics across the lifespan (Zappasodi et al., 2015). Within this framework, the enhanced interhemispheric propagation observed in our study may reflect a reorganization toward less segregated and more distributed communication patterns in the aging brain. Such changes could represent compensatory adaptations that support functional integration despite declining local specialization, although interpretations in terms of neural dedifferentiation or reduced network specificity also remain plausible.

The present findings also contribute to the broader debate regarding the functional role of alpha oscillations in aging. Alpha activity has long been associated with inhibitory control, sensory gating, attentional allocation, predictive processing, and large-scale coordination of cortical processing (Klimesch et al., 2007b; Mathewson et al., 2011; Klimesch, 2012; Lobier et al., 2018; Schneider et al., 2019; Tarasi et al., 2022, 2023, 2026a). Several task-based studies indicate that aging impairs the ability to dynamically modulate alpha rhythms in response to behavioral demands. For example, younger adults typically exhibit alpha synchronization during motor inhibition and alpha desynchronization during movement execution, whereas these modulations are markedly reduced or absent in older adults (Bönstrup et al., 2015; Manor et al., 2023). Similarly, older adults appear less able to regulate alpha power to suppress distracting information (Vaden et al., 2012), suggesting age-related alterations in inhibitory and selective-attention mechanisms. At the same time, some studies have reported enhanced frontal alpha phase-locking or increased frontal alpha responses in middle-aged and older individuals (Yordanova et al., 1998; Kolev et al., 2002), supporting the idea that aging involves a redistribution rather than a simple loss of alpha activity. This interpretation aligns with broader models of functional reorganization, including HAROLD and PASA, in which older adults recruit more distributed or frontal networks to sustain cognitive performance. Importantly, the relationship between alpha dynamics and cognition in aging remains complex. While some studies found no direct association between resting-state alpha power or peak alpha frequency and cognitive performance (Finnigan and Robertson, 2011), others suggest that reductions in alpha power and slowing of alpha peak frequency may serve as markers of cognitive decline and mnemonic dysfunction (Lejko et al., 2020; Puttaert et al., 2021). Notably, one study demonstrated that when task performance is equated between younger and older adults, alpha power and frequency can still be modulated similarly across working-memory loads, suggesting that preserved alpha dynamics may support maintained cognitive function in aging (Sghirripa et al., 2021). In this context, our findings of altered alpha power and connectivity may reflect adaptive network-level reorganization rather than purely pathological decline, which has been shown to relate to aging and lack of impaired cognitive functions (Trammell et al., 2017).

Taken together, our findings suggest that aging is characterized by a complex reorganization of alpha-band dynamics involving reduced oscillatory power, altered large-scale connectivity, and increased interhemispheric propagation in frontal regions. Future studies employing attentionally demanding or visually driven paradigms may help clarify how these resting-state alterations influence the functional role of alpha oscillations during active cognitive processing.

## Conflict of interest statement

The authors declare no competing financial interests.

## Author contribution statement

AA, LT, and VR conceived the idea. LT and VR collected the data. PMM contributed to the conceptualization and literature overview. AA, JS, and LT analyzed the data. AA, VR, and LT acquired financial support. AA wrote the first draft and figure visualization. All authors revised and contributed to the final version.

## Acknowledgments

The authors thank the participants for their time. A.A. was funded by the European Union under the European Union’s Horizon 2020 research and innovation program (grant agreement No. 101075930). The copyright holder for this is of the author(s) only and does not necessarily reflect those of the European Union or the European Research Council (ERC). Neither the European Union nor the granting authority can be held responsible for them.V.R. is supported by Next Generation EU (NGEU) and funded by the Ministry of the University and Research (MUR), National Recovery and Research Plan (NRRP) PRIN 2022 (grant n 2022H4ZRSN—CUP J53D23008040006): predictive waves in human perception and individual differences along the autism-schizophrenia continuum (D DN. 104 02.02.2022); (grant n. P2022XAKXL—CUP J53D23017340001): Investigating the plasticity of human predictive coding through neuromodulation (D DN. 1409 14.09.2022); Ministerio de Ciencia, Innovación y Universidades, Spain (PID2019-111335 GA-100); L.T. is supported by Bial Foundation (219/24).

## Supplementary materials

**Figure S1:**
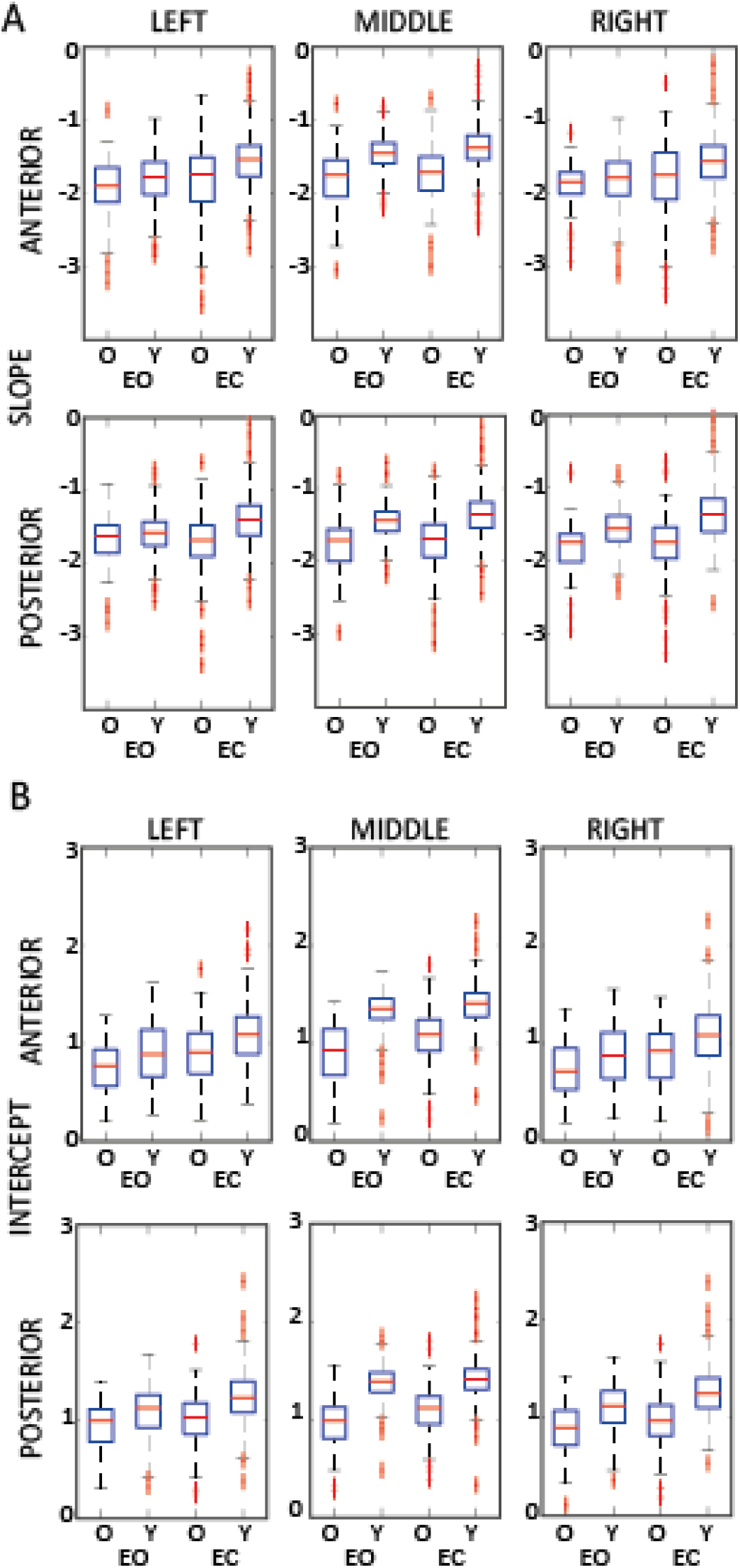
Group differences in aperiodic spectral parameters during resting-state EEG divided per regions. Boxplots show the average values across all electrodes for the aperiodic **slope** (A) and **intercept** (B) in older (O) and younger (Y) adults during eyes-open (EO) and eyes-closed (EC) resting-state recordings. Boxes indicate the interquartile range (IQR), center lines denote the median, whiskers extend to 1.5 × IQR, and red dots represent individual participants. The columns represent the left, middle and right areas in the anterior (first row) and posterior (second row) regions, respectively.

**Figure S2:**
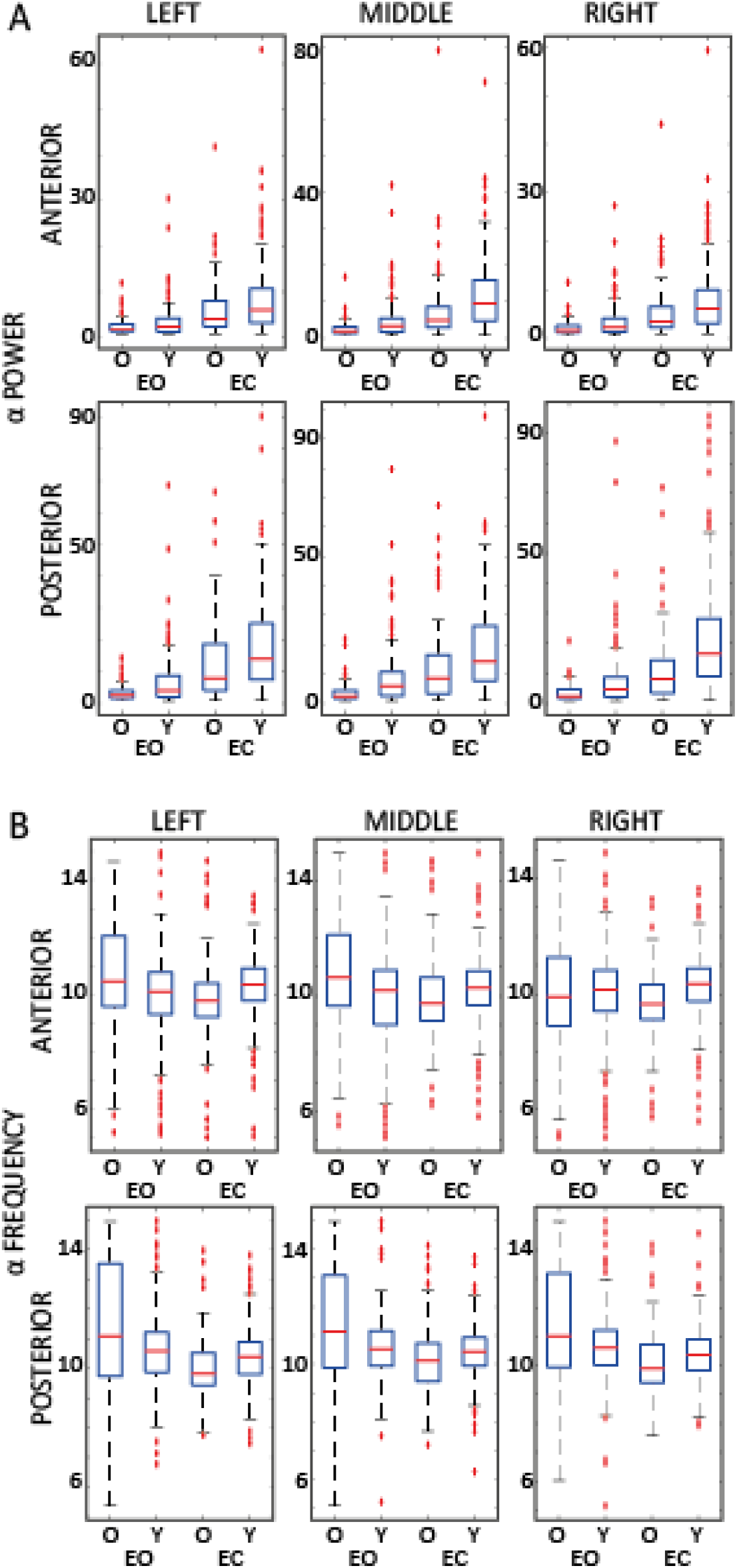
Group differences in α-power and α peak frequency during resting-state EEG divided per regions. Boxplots show the average values across all electrodes for the aperiodic **power** (A) and **peak frequency** (B) in older (O) and younger (Y) adults during eyes-open (EO) and eyes-closed (EC) resting-state recordings. Boxes indicate the interquartile range (IQR), center lines denote the median, whiskers extend to 1.5 × IQR, and red dots represent individual participants. The columns represent the left, middle and right areas in the anterior (first row) and posterior (second row) regions, respectively.

**Figure S3:**
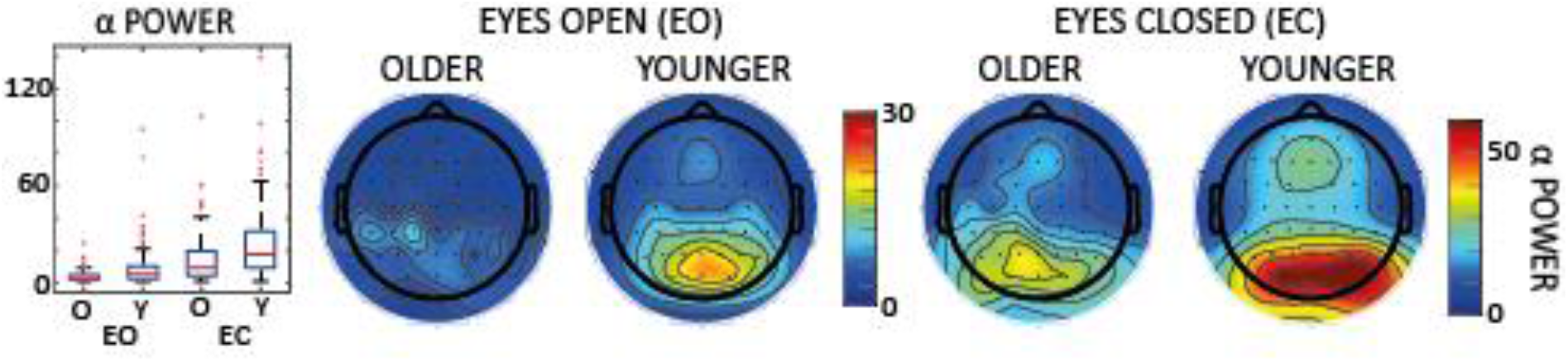
Group differences in α-power centered around the α peak frequency. Boxplots show the average values across all electrodes for the alpha-band power in older (O) and younger (Y) adults during eyes-open (EO) and eyes-closed (EC) resting-state recordings. Boxes indicate the interquartile range (IQR), center lines denote the median, whiskers extend to 1.5 × IQR, and red dots represent individual participants. Scalp topographies show the spatial distribution for younger and older adults during EO and EC. Warmer colors indicate higher power and cooler colors indicate lower values. Contour lines represent interpolated isovalues across the scalp. Topographic maps are displayed with the nose at the top and the left hemisphere on the left.

## Notes

### Competing Interest Statement

The authors have declared no competing interest.

## References

Alamia A, Terral L, d’Ambra MR, VanRullen R (2023) Distinct roles of forward and backward alpha-band waves in spatial visual attention. Elife 12:e85035.

Alamia A, VanRullen R (2019) Alpha oscillations and traveling waves: Signatures of predictive coding? PLOS Biology 17:e3000487.

Alamia A, VanRullen R (2023) A Traveling Waves Perspective on Temporal Binding. J Cogn Neurosci:1–9.

Babiloni C, Binetti G, Cassarino A, Dal Forno G, Del Percio C, Ferreri F, Ferri R, Frisoni G, Galderisi S, Hirata K, Lanuzza B, Miniussi C, Mucci A, Nobili F, Rodriguez G, Luca Romani G, Rossini PM (2006) Sources of cortical rhythms in adults during physiological aging: a multicentric EEG study. Hum Brain Mapp 27:162–172.

Balsters JH, O’Connell RG, Galli A, Nolan H, Greco E, Kilcullen SM, Bokde ALW, Lai R, Upton N, Robertson IH (2013) Changes in resting connectivity with age: a simultaneous electroencephalogram and functional magnetic resonance imaging investigation. Neurobiology of Aging 34:2194–2207.

Bastos AM, Vezoli J, Bosman CA, Schoffelen J-M, Oostenveld R, Dowdall JR, De Weerd P, Kennedy H, Fries P (2015) Visual Areas Exert Feedforward and Feedback Influences through Distinct Frequency Channels. Neuron 85:390–401.

Bauer M, Stenner M-P, Friston KJ, Dolan RJ (2014) Attentional Modulation of Alpha/Beta and Gamma Oscillations Reflect Functionally Distinct Processes. J Neurosci 34:16117–16125.

Berlingeri M, Danelli L, Bottini G, Sberna M, Paulesu E (2013) Reassessing the HAROLD model: Is the hemispheric asymmetry reduction in older adults a special case of compensatory-related utilisation of neural circuits? Exp Brain Res 224:393–410.

Bonnefond M, Jensen O (2012) Alpha Oscillations Serve to Protect Working Memory Maintenance against Anticipated Distracters. Current Biology 22:1969–1974.

Bönstrup M, Hagemann J, Gerloff C, Sauseng P, Hummel FC (2015) Alpha oscillatory correlates of motor inhibition in the aged brain. Front Aging Neurosci 7:193.

Bressler SL, Richter CG (2015) Interareal oscillatory synchronization in top-down neocortical processing. Current Opinion in Neurobiology 31:62–66.

Brüers S, VanRullen R (2018) Alpha Power Modulates Perception Independently of Endogenous Factors. Front Neurosci 12 Available at: https://www.frontiersin.org/journals/neuroscience/articles/10.3389/fnins.2018.00279/full [Accessed May 9, 2026].

Busch NA, Dubois J, VanRullen R (2009) The Phase of Ongoing EEG Oscillations Predicts Visual Perception. J Neurosci 29:7869–7876.

Buzsáki G, Draguhn A (2004) Neuronal Oscillations in Cortical Networks. Science 304:1926–1929.

Cabeza R (2002) Hemispheric asymmetry reduction in older adults: the HAROLD model. Psychol Aging 17:85–100.

Chiang AKI, Rennie CJ, Robinson PA, van Albada SJ, Kerr CC (2011) Age trends and sex differences of alpha rhythms including split alpha peaks. Clinical Neurophysiology 122:1505–1517.

Damoiseaux JS (2017) Effects of aging on functional and structural brain connectivity. NeuroImage 160:32–40.

Das A, Zabeh E, Jacobs J (2022) How can we detect and analyze traveling waves in human brain oscillations? Available at: https://osf.io/jhnpr_v1 [Accessed May 7, 2025].

Davis SW, Dennis NA, Daselaar SM, Fleck MS, Cabeza R (2008) Que PASA? The posterior-anterior shift in aging. Cereb Cortex 18:1201–1209.

Deery HA, Di Paolo R, Moran C, Egan GF, Jamadar SD (2023) The older adult brain is less modular, more integrated, and less efficient at rest: A systematic review of large-scale resting-state functional brain networks in aging. Psychophysiology 60:e14159.

Dolcos F, Rice HJ, Cabeza R (2002) Hemispheric asymmetry and aging: right hemisphere decline or asymmetry reduction. Neuroscience & Biobehavioral Reviews 26:819–825.

Donoghue T, Haller M, Peterson EJ, Varma P, Sebastian P, Gao R, Noto T, Lara AH, Wallis JD, Knight RT, Shestyuk A, Voytek B (2020) Parameterizing neural power spectra into periodic and aperiodic components. Nat Neurosci 23:1655–1665.

Dushanova J, Christov M (2014) The effect of aging on EEG brain oscillations related to sensory and sensorimotor functions. Advances in Medical Sciences 59:61–67.

Fakche C, VanRullen R, Marque P, Dugué L (2022) α Phase-Amplitude Tradeoffs Predict Visual Perception. eNeuro 9 Available at: https://www.eneuro.org/content/9/1/ENEURO.0244-21.2022 [Accessed December 29, 2024].

Finnigan S, Robertson IH (2011) Resting EEG theta power correlates with cognitive performance in healthy older adults. Psychophysiology 48:1083–1087.

Frisoni M, Tarasi L, Borgomaneri S, Romei V (2025) The relationship between individual alpha frequency and time perception: Testing the internal clock versus the sampling rate hypothesis. Cortex 192:183–195.

Gaál ZA, Boha R, Stam CJ, Molnár M (2010) Age-dependent features of EEG-reactivity—Spectral, complexity, and network characteristics. Neuroscience Letters 479:79–84.

Gómez C, Pérez-Macías JM, Poza J, Fernández A, Hornero R (2013) Spectral changes in spontaneous MEG activity across the lifespan. J Neural Eng 10:066006.

Grandy TH, Werkle-Bergner M, Chicherio C, Schmiedek F, Lövdén M, Lindenberger U (2013) Peak individual alpha frequency qualifies as a stable neurophysiological trait marker in healthy younger and older adults. Psychophysiology 50:570–582.

Haigh ZJ, Tran H, Berger T, Shirinpour S, Alekseichuk I, Koenig S, Zimmermann J, McGovern R, Darrow D, Herman A, Wischnewski M, Opitz A (2025) Modulation of motor excitability reflects traveling waves of neural oscillations. Cell Reports 44 Available at: https://www.cell.com/cell-reports/abstract/S2211-1247(25)00635-7 [Accessed May 9, 2026].

Händel BF, Haarmeier T, Jensen O (2011) Alpha Oscillations Correlate with the Successful Inhibition of Unattended Stimuli. J Cogn Neurosci 23:2494–2502.

He X, Ruan X, Shen M, Yuan J, Li C, Yang Y, Zhu J, Cui R, Lu Z-L, Chen J-F, Hou F (2025) The temporal window of visual processing throughout adulthood. Front Neurosci 19 Available at: https://www.frontiersin.org/journals/neuroscience/articles/10.3389/fnins.2025.1547959/full [Accessed May 20, 2026].

He X, Shen M, Cui R, Zheng H, Ruan X, Lu Z-L, Hou F (2020) The Temporal Window of Visual Processing in Aging. Invest Ophthalmol Vis Sci 61:60.

Ishii R, Canuet L, Aoki Y, Hata M, Iwase M, Ikeda S, Nishida K, Ikeda M (2017) Healthy and Pathological Brain Aging: From the Perspective of Oscillations, Functional Connectivity, and Signal Complexity. Neuropsychobiology 75:151–161.

Jensen O, Mazaheri A (2010) Shaping Functional Architecture by Oscillatory Alpha Activity: Gating by Inhibition. Front Hum Neurosci 4 Available at: https://www.frontiersin.org/journals/human-neuroscience/articles/10.3389/fnhum.2010.00186/full [Accessed May 9, 2026].

Keitel C, Keitel A, Benwell CSY, Daube C, Thut G, Gross J (2019) Stimulus-Driven Brain Rhythms within the Alpha Band: The Attentional-Modulation Conundrum. J Neurosci 39:3119–3129.

Kikuchi M, Wada Y, Koshino Y, Nanbu Y, Hashimoto T (2000) Effect of Normal Aging upon Interhemispheric EEG Coherence: Analysis during Rest and Photic Stimulation. Clinical Electroencephalography 31:170–174.

Klimesch W (1997) EEG-alpha rhythms and memory processes. International Journal of Psychophysiology 26:319–340.

Klimesch W (1999) EEG alpha and theta oscillations reflect cognitive and memory performance: a review and analysis. Brain Research Reviews 29:169–195.

Klimesch W (2012) Alpha-band oscillations, attention, and controlled access to stored information. Trends in Cognitive Sciences 16:606–617.

Klimesch W, Hanslmayr S, Sauseng P, Gruber WR, Doppelmayr M (2007a) P1 and Traveling Alpha Waves: Evidence for Evoked Oscillations. Journal of Neurophysiology 97:1311–1318.

Klimesch W, Sauseng P, Hanslmayr S (2007b) EEG alpha oscillations: The inhibition–timing hypothesis. Brain Research Reviews 53:63–88.

Knyazeva MG, Barzegaran E, Vildavski VY, Demonet J-F (2018) Aging of human alpha rhythm. Neurobiology of Aging 69:261–273.

Kolev V, Yordanova J, Basar-Eroglu C, Basar E (2002) Age effects on visual EEG responses reveal distinct frontal alpha networks. Clinical Neurophysiology 113:901–910.

Lejko N, Larabi DI, Herrmann CS, Aleman A, Ćurčić-Blake B (2020) Alpha Power and Functional Connectivity in Cognitive Decline: A Systematic Review and Meta-Analysis. Journal of Alzheimer’s Disease 78:1047–1088.

Li H-J, Hou X-H, Liu H-H, Yue C-L, Lu G-M, Zuo X-N (2015) Putting age-related task activation into large-scale brain networks: A meta-analysis of 114 fMRI studies on healthy aging. Neuroscience & Biobehavioral Reviews 57:156–174.

Lobier M, Palva JM, Palva S (2018) High-alpha band synchronization across frontal, parietal and visual cortex mediates behavioral and neuronal effects of visuospatial attention. Neuroimage 165:222–237.

Love J, Selker R, Marsman M, Jamil T, Dropmann D, Verhagen J, Ly A, Gronau QF, Šmíra M, Epskamp S, Matzke D, Wild A, Knight P, Rouder JN, Morey RD, Wagenmakers E-J (2019) JASP: Graphical Statistical Software for Common Statistical Designs. Journal of Statistical Software 88:1–17.

Lozano-Soldevilla D, VanRullen R (2019) The Hidden Spatial Dimension of Alpha: 10-Hz Perceptual Echoes Propagate as Periodic Traveling Waves in the Human Brain. Cell Reports 26:374–380.e4.

Luo C, Ester EF (2025) Traveling waves link human visual and frontal cortex during working memory– guided behavior. Proceedings of the National Academy of Sciences 122:e2415573122.

Manor R, Cheaha D, Kumarnsit E, Samerphob N (2023) Age-related Deterioration of Alpha Power in Cortical Areas Slowing Motor Command Formation in Healthy Elderly Subjects. In Vivo 37:679–684.

Mathewson KE, Lleras A, Beck DM, Fabiani M, Ro T, Gratton G (2011) Pulsed Out of Awareness: EEG Alpha Oscillations Represent a Pulsed-Inhibition of Ongoing Cortical Processing. Front Psychol 2 Available at: https://www.frontiersin.org/journals/psychology/articles/10.3389/fpsyg.2011.00099/full [Accessed May 15, 2026].

Mayer A, Schwiedrzik CM, Wibral M, Singer W, Melloni L (2016) Expecting to See a Letter: Alpha Oscillations as Carriers of Top-Down Sensory Predictions. Cereb Cortex 26:3146–3160.

Mccarthy P, Benuskova L, Franz EA (2014) The age-related posterior-anterior shift as revealed by voxelwise analysis of functional brain networks. Frontiers in Aging Neuroscience Available at: https://www.proquest.com/docview/2301923682/abstract/44D18479539D49DBPQ/1 [Accessed May 15, 2026].

Merkin A, Sghirripa S, Graetz L, Smith AE, Hordacre B, Harris R, Pitcher J, Semmler J, Rogasch NC, Goldsworthy M (2023) Do age-related differences in aperiodic neural activity explain differences in resting EEG alpha? Neurobiol Aging 121:78–87.

Michalareas G, Vezoli J, van Pelt S, Schoffelen J-M, Kennedy H, Fries P (2016) Alpha-Beta and Gamma Rhythms Subserve Feedback and Feedforward Influences among Human Visual Cortical Areas. Neuron 89:384–397.

Mohan UR, Zhang H, Ermentrout B, Jacobs J (2024) The direction of theta and alpha travelling waves modulates human memory processing. Nat Hum Behav 8:1124–1135.

Muller L, Chavane F, Reynolds J, Sejnowski TJ (2018) Cortical travelling waves: mechanisms and computational principles. Nat Rev Neurosci 19:255–268.

Palva S, Palva JM (2007) New vistas for α-frequency band oscillations. Trends in Neurosciences 30:150–158.

Pang (庞兆阳) Z, Alamia A, VanRullen R (2020) Turning the Stimulus On and Off Changes the Direction of α Traveling Waves. eNeuro 7:ENEURO.0218-20.2020.

Park J, Ho RLM, Wang W-E, Nguyen VQ, Coombes SA (2024) The effect of age on alpha rhythms in the human brain derived from source localized resting-state electroencephalography. Neuroimage 292:120614.

Perinelli A, Assecondi S, Tagliabue CF, Mazza V (2022) Power shift and connectivity changes in healthy aging during resting-state EEG. Neuroimage 256:119247.

Polich J (1997) EEG and ERP assessment of normal aging. Electroencephalogr Clin Neurophysiol 104:244–256.

Puttaert D, Wens V, Fery P, Rovai A, Trotta N, Coquelet N, De Breucker S, Sadeghi N, Coolen T, Goldman S, Peigneux P, Bier J-C, De Tiège X (2021) Decreased Alpha Peak Frequency Is Linked to Episodic Memory Impairment in Pathological Aging. Front Aging Neurosci 13:711375.

Reuter-Lorenz PA, Cappell KA (2008) Neurocognitive Aging and the Compensation Hypothesis. Curr Dir Psychol Sci 17:177–182.

Reuter-Lorenz PA, Park DC (2014) How Does it STAC Up? Revisiting the Scaffolding Theory of Aging and Cognition. Neuropsychol Rev 24:355–370.

Rohenkohl G, Nobre AC (2011) Alpha Oscillations Related to Anticipatory Attention Follow Temporal Expectations. J Neurosci 31:14076–14084.

Romei V, Tarasi L (2026) Alpha frequency shapes perceptual sensitivity by modulating optimal phase likelihood. Nat Commun 17:3384.

Ruzzoli M, Torralba M, Morís Fernández L, Soto-Faraco S (2019) The relevance of alpha phase in human perception. Cortex 120:249–268.

Sadaghiani S, Kleinschmidt A (2016) Brain Networks and α-Oscillations: Structural and Functional Foundations of Cognitive Control. Trends in Cognitive Sciences 20:805–817.

Sala-Llonch R, Bartrés-Faz D, Junqué C (2015) Reorganization of brain networks in aging: a review of functional connectivity studies. Front Psychol 6 Available at: https://www.frontiersin.org/journals/psychology/articles/10.3389/fpsyg.2015.00663/full [Accessed July 20, 2026].

Samaha J, Romei V (2024) Alpha-Band Frequency and Temporal Windows in Perception: A Review and Living Meta-analysis of 27 Experiments (and Counting). J Cogn Neurosci 36:640–654.

Sato TK, Nauhaus I, Carandini M (2012) Traveling Waves in Visual Cortex. Neuron 75:218–229.

Sauseng P, Klimesch W, Stadler W, Schabus M, Doppelmayr M, Hanslmayr S, Gruber WR, Birbaumer N (2005) A shift of visual spatial attention is selectively associated with human EEG alpha activity. European Journal of Neuroscience 22:2917–2926.

Scally B, Burke MR, Bunce D, Delvenne J-F (2018) Resting-state EEG power and connectivity are associated with alpha peak frequency slowing in healthy aging. Neurobiol Aging 71:149–155.

Schneider D, Göddertz A, Haase H, Hickey C, Wascher E (2019) Hemispheric asymmetries in EEG alpha oscillations indicate active inhibition during attentional orienting within working memory. Behavioural Brain Research 359:38–46.

Schwenk JCB, Alamia A (2025) Detection and quantification of planar traveling waves in the EEG using spherical phase fitting.:2025.12.03.692197 Available at: https://www.biorxiv.org/content/10.64898/2025.12.03.692197v1 [Accessed December 21, 2025].

Sghirripa S, Graetz L, Merkin A, Rogasch NC, Semmler JG, Goldsworthy MR (2021) Load-dependent modulation of alpha oscillations during working memory encoding and retention in young and older adults. Psychophysiology 58:e13719.

Song J, Birn RM, Boly M, Meier TB, Nair VA, Meyerand ME, Prabhakaran V (2014) Age-related reorganizational changes in modularity and functional connectivity of human brain networks. Brain Connect 4:662–676.

Spreng RN, Turner GR (2019) The Shifting Architecture of Cognition and Brain Function in Older Adulthood. Perspect Psychol Sci 14:523–542.

Stacey JE, Crook-Rumsey M, Sumich A, Howard CJ, Crawford T, Livne K, Lenzoni S, Badham S (2021) Age differences in resting state EEG and their relation to eye movements and cognitive performance. Neuropsychologia 157:107887.

Tarasi L, Alamia A, Romei V (2025a) Perceptual Bias in Motion Discrimination is Related to Asymmetric Interhemispheric Alpha Traveling Waves. Advanced Science 12:e14623.

Tarasi L, Alamia A, Romei V (2026a) Backward alpha band oscillations shape perceptual bias under probabilistic cues. Commun Biol 9:280.

Tarasi L, Bertaccini R, Ippolito G, Martelli ME, di Pellegrino G, Romei V (2025b) Oscillatory signatures of monitoring and anticipatory strategies for probabilistic vs deterministic cues. Imaging Neuroscience 3:imag_a_00496.

Tarasi L, Covelli M, Fatis CT de, Bertini C, Avenanti A, Romei V (2026b) Aging redirects prior expectations from perception to action.:2026.01.09.698574 Available at: https://www.biorxiv.org/content/10.64898/2026.01.09.698574v1 [Accessed May 18, 2026].

Tarasi L, di Pellegrino G, Romei V (2022) Are you an empiricist or a believer? Neural signatures of predictive strategies in humans. Progress in Neurobiology 219:102367.

Tarasi L, Martelli ME, Bortoletto M, di Pellegrino G, Romei V (2023) Neural Signatures of Predictive Strategies Track Individuals Along the Autism-Schizophrenia Continuum. Schizophr Bull 49:1294–1304.

Tarasi L, Romanazzi D, Pasini A, Romei V (2025c) Delusion-like thinking is associated with lower individual alpha peak frequency. Schizophr 11:76.

Tarasi L, Romei V (2024) Individual Alpha Frequency Contributes to the Precision of Human Visual Processing. J Cogn Neurosci 36:602–613.

Thut G, Nietzel A, Brandt SA, Pascual-Leone A (2006) α-Band Electroencephalographic Activity over Occipital Cortex Indexes Visuospatial Attention Bias and Predicts Visual Target Detection. J Neurosci 26:9494–9502.

Tomasi D, Volkow ND (2012) Aging and functional brain networks. Mol Psychiatry 17:549–558.

Trajkovic J, Ricci G, Pirazzini G, Tarasi L, Di Gregorio F, Magosso E, Ursino M, Romei V (2025a) Aberrant Functional Connectivity and Brain Network Organization in High-Schizotypy Individuals: An Electroencephalography Study. Schizophr Bull 51:1266–1281.

Trajkovic J, Veniero D, Hanslmayr S, Palva S, Cruz G, Romei V, Thut G (2025b) Top-down and bottom-up interactions rely on nested brain oscillations to shape rhythmic visual attention sampling. PLOS Biology 23:e3002688.

Trammell JP, MacRae PG, Davis G, Bergstedt D, Anderson AE (2017) The Relationship of Cognitive Performance and the Theta-Alpha Power Ratio Is Age-Dependent: An EEG Study of Short Term Memory and Reasoning during Task and Resting-State in Healthy Young and Old Adults. Front Aging Neurosci 9 Available at: https://www.frontiersin.org/journals/aging-neuroscience/articles/10.3389/fnagi.2017.00364/full [Accessed December 11, 2025].

Tröndle M, Popov T, Pedroni A, Pfeiffer C, Barańczuk-Turska Z, Langer N (2023) Decomposing age effects in EEG alpha power. Cortex 161:116–144.

Ursino M, Serra M, Tarasi L, Ricci G, Magosso E, Romei V (2022) Bottom-up vs. top-down connectivity imbalance in individuals with high-autistic traits: An electroencephalographic study. Front Syst Neurosci 16 Available at: https://www.frontiersin.org/journals/systems-neuroscience/articles/10.3389/fnsys.2022.932128/full [Accessed April 15, 2026].

Vaden RJ, Hutcheson NL, McCollum LA, Kentros JG, Visscher KM (2012) Older adults, unlike younger adults, do not modulate alpha power to suppress irrelevant information. Neuroimage 63:1127–1133.

van Diepen RM, Cohen MX, Denys D, Mazaheri A (2015) Attention and Temporal Expectations Modulate Power, Not Phase, of Ongoing Alpha Oscillations. J Cogn Neurosci 27:1573–1586.

VanRullen R (2016) Perceptual Cycles. Trends in Cognitive Sciences 20:723–735.

VanRullen R, Macdonald JSP (2012) Perceptual Echoes at 10 Hz in the Human Brain. Current Biology 22:995–999.

Vecchio F, Miraglia F, Bramanti P, Rossini PM (2014) Human Brain Networks in Physiological Aging: A Graph Theoretical Analysis of Cortical Connectivity from EEG Data. Journal of Alzheimer’s Disease 41:1239–1249.

Vysata O, Kukal J, Prochazka A, Pazdera L, Simko J, Valis M (2014) Age-related changes in EEG coherence. Neurologia i Neurochirurgia Polska 48:35–38.

Ward LM (2003) Synchronous neural oscillations and cognitive processes. Trends in Cognitive Sciences 7:553–559.

Wei J, Alamia A, Yao Z, Huang G, Li L, Liang Z, Zhang L, Zhou C, Song Z, Zhang Z (2024) State-Dependent tACS Effects Reveal the Potential Causal Role of Prestimulus Alpha Traveling Waves in Visual Contrast Detection. J Neurosci 44 Available at: https://www.jneurosci.org/content/44/27/e2023232024 [Accessed May 7, 2025].

Wianda E, Ross B (2019) The roles of alpha oscillation in working memory retention. Brain and Behavior 9:e01263.

Worden MS, Foxe JJ, Wang N, Simpson GV (2000) Anticipatory Biasing of Visuospatial Attention Indexed by Retinotopically Specific α-Bank Electroencephalography Increases over Occipital Cortex. J Neurosci 20:RC63–RC63.

Yordanova JY, Kolev VN, Başar E (1998) EEG theta and frontal alpha oscillations during auditory processing change with aging. Electroencephalography and Clinical Neurophysiology/Evoked Potentials Section 108:497–505.

Zappasodi F, Marzetti L, Olejarczyk E, Tecchio F, Pizzella V (2015) Age-Related Changes in Electroencephalographic Signal Complexity. PLoS One 10:e0141995.

Zeng Y, Sauseng P, Alamia A (2024) Alpha Traveling Waves during Working Memory: Disentangling Bottom-Up Gating and Top-Down Gain Control. J Neurosci 44 Available at: https://www.jneurosci.org/content/44/50/e0532242024 [Accessed March 18, 2025].

Zhang H, Watrous AJ, Patel A, Jacobs J (2018) Theta and Alpha Oscillations Are Traveling Waves in the Human Neocortex. Neuron 98:1269–1281.e4.

Zich C, Quinn AJ, Bonaiuto JJ, O’Neill G, Mardell LC, Ward NS, Bestmann S (2023) Spatiotemporal organisation of human sensorimotor beta burst activity Vinck M, Baker CI, Law R, eds. eLife 12:e80160.

